# Antimalarial drugs lose their activity with a slight drop in pH

**DOI:** 10.1101/2020.08.04.237255

**Authors:** Tomohisa Kitagawa, Atsushi Mastumoto, Ichiro Terashima, Yukifumi Uesono

## Abstract

Antimalarial drugs have antimicrobial, antiviral, antimalarial and immunosuppressive activities, although the mechanisms remain unknown. Quinacrine (QC) increases the antimicrobial activity against yeast exponentially with a pH-dependent increase in the cationic amphiphilic drug (CAD) structure. CAD-QC localizes in membranes and induces glucose starvation by noncompetitively inhibiting glucose uptake. A logarithmic increase in antimicrobial activity with pH-dependent CAD formation was also observed for chloroquine, indicating that the CAD structure is crucial for its pharmacological activity. A decrease in CAD structure with a slight decrease in pH from 7.4 greatly reduced their effects; namely, these drugs would inefficiently act on falciparum malaria and COVID-19 pneumonia patients with acidosis, resulting in resistance. Recovering normal blood pH or using pH-insensitive quinoline drugs might be effective.

## Introduction

Malaria is one of the most serious infectious parasitic diseases in the world. Quinine (QN) was the first natural compound to successfully treat malaria. Thereafter, quinacrine (QC) was developed in 1932 as a synthetic antimalarial drug as an alternative to quinine. QC was used during World War II until 1945 when it was substituted by chloroquine (CQ) (*1*). Currently, CQ is used as an antimalarial drug, although it has no effect on falciparum malaria, and the resistance mechanism has not been fully elucidated (*2*, *3*). Additionally, QC, CQ and hydroxychloroquine (HCQ) have been clinically used as immunosuppressive agents for refractory lupus erythematosus and rheumatoid arthritis (*2*). These antimalarial drugs have various pharmacological effects, such as antiviral, antimicrobial, and anticancer activities at the cellular level, and thrombosis prevention (*2*, *4–15*). Therefore, these drugs have the potential to be useful agents without any significant clinical concerns because of a long history of clinical use. However, their cellular effects have not often been confirmed at the whole-body level (*16*). For drug development, it is necessary to elucidate the discrepancy. Various action mechanisms for antimalarial drugs have been proposed. CQ accumulates in acidic food vacuoles of the malaria parasite, thereby exerting antimalarial effects by increasing the pH or accumulating toxic heme in the vacuoles (*2*, *17*, *18*). A similar vacuole accumulation mechanism has been considered for QC (*2*, *19*). Additionally, QC is thought to inhibit DNA/RNA synthesis (*20*), calmodulin-mediated signaling (*21*), electron transport (*22*), and phospholipase A2, and interact with membrane phospholipids (*23*, *24*). However, there is currently no unified view to explain the pleiotropic effects of QC and CQ analogs. Several antimalarial drugs inhibit the growth of pathogenic and budding yeasts without hemoglobin (*7*–*14*). Because QC accumulates in the acidic vacuoles of yeast but not in the nuclei (*25*), this might be a clue to elucidate the antimicrobial mechanism.

Antipsychotic phenothiazines, local anesthetics, and cationic surfactants are cationic amphiphilic drugs (CADs) that induce the glucose starvation mimetic state in budding yeast irrespective of their structural differences (*26*). Thus, the CAD structure is important to elucidate the common mechanism. Chlorpromazine (CPZ), a phenothiazine, is the first synthetic compound with antipsychotic effects (*27*), and its structure is similar to that of QC. Thus, QC might also be CAD and induce CAD-like effects in yeast. Here, we demonstrate both the pH-dependent structural changes in QC and the nonspecific interactions of the CAD structure with the membrane are crucial for antimicrobial activity. Additionally, we describe the mechanism by which QC inhibits hexose transporter function as a target. Finally, we describe the importance of the CAD structure common to antimalarial quinoline drugs on resistance and propose effective clinical uses of these drugs.

## Results

### Structures and physicochemical properties of QC change drastically in response to pH

The tertiary propylamine tail of CPZ is hydrophilic, and the tricyclic ring (phenothiazine) structure is hydrophobic, thereby forming the CAD structure. Thus, it binds directly to artificial membranes without proteins (*28*) and is localized to yeast membranes (*26*, *29*) (Fig. 1A). QC is also considered to form a CAD structure that contains butyl amino groups as the hydrophilic region and acridine rings as the hydrophobic region. However, QC localizes to the acidic vacuoles of yeast, suggesting that QC is not a simple CAD. To ensure the CAD structures of these compounds, we estimated the physicochemical properties using computational analysis. The analysis estimated that QC had two acid dissociation constants (pKa 1: 8.37, pKa 2: 10.33) (Fig. 1B). Because the pKa values of QC are attributed to protonation of the nitrogen in the acridine ring (a) and of the side chain (b), QC forms three structures in response to pH (Fig. 1B). The nitrogen atom positively charged by protonation interacts with the negatively charged oxygen of water, thereby increasing water solubility. Thus, the major structures of QC were the hydrophilic form (HP-QC) at pH <8.37, the CAD form (CAD-QC) at pH 8.37-10.33, and the lipophilic form (LP-QC) at pH >10.33. The software also estimated the logD, the pH-dependent octanol/water partition coefficient (logP) reflecting the lipophilicity of drugs. These analyses indicate that the lipophilicity of QC increases with increasing CAD-QC at pH values greater than 6. Amphiphilic compounds form micelles, a characteristic property of surfactants, at high concentrations. Because the critical micelle concentration of QC is 70 mM in aqueous solution (*30*), QC would form an amphiphilic structure at neutral pH. Similar pH-dependent structural changes were estimated in CQ with two pKa values (7.29 and 10.32) (Fig. S1). In contrast, CPZ has a pKa of 9.2, reflecting protonation of only the nitrogen of the side chain. Thus, the major structures of CPZ are CAD-CPZ at pH <9.2 and LP-CPZ at pH >9.2 (Fig. S2). The lipophilicity of CPZ increased with increasing LP-CPZ, although the degree of increase was smaller than those of QC and CQ, as judged by the logD. These estimations suggest that the lipophilicities of the antimalarial drugs change drastically in response to pH-dependent structural change compared with CPZ.

**Fig. 1.**
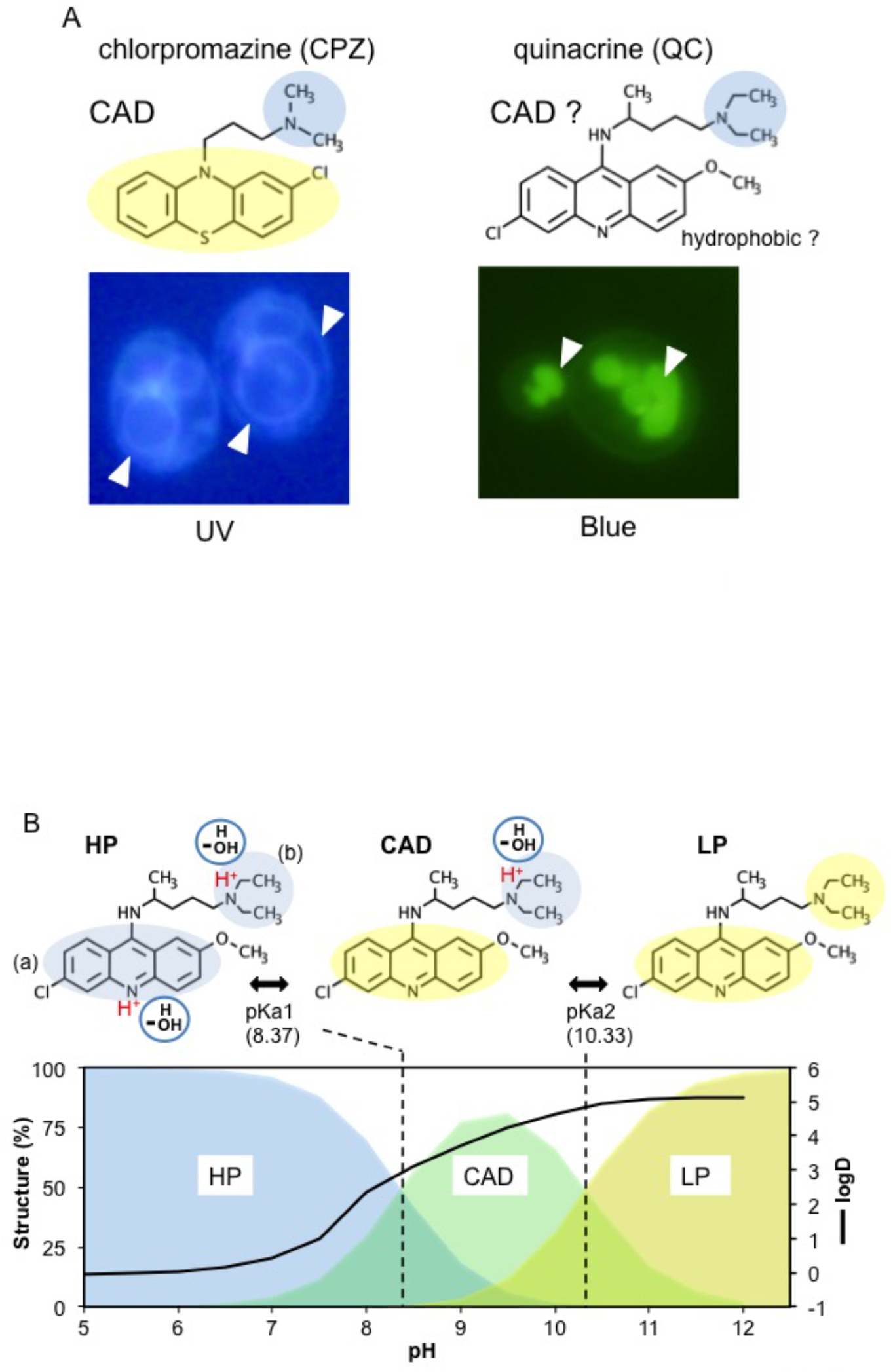
Structures and physicochemical properties of quinacrine. **(A)** Comparison between chlorpromazine and quinacrine. Upper panels: Structures of chlorpromazine (CPZ) (left) and quinacrine (QC) (right). Blue and yellow areas indicate hydrophilic and hydrophobic regions, respectively. Lower panels: BY4741 cells treated with 100 mM phosphate buffer (pH 7.4) containing 2% glucose and stained with 50 μM CPZ and 10 μM QC for 30 min at 25°C. The fluorescence signals for CPZ and QC excited by UV and blue light irradiation are shown. **(B)** Estimated change in the structural and physicochemical properties of QC as a function of pH. The structures and physicochemical properties were estimated by MarvinSketch software. Upper panel: The pH-dependent QC structures are shown as the hydrophilic (HP), cationic amphiphilic (CAD), and lipophilic (LP) forms. Blue and yellow areas indicate hydrophilic and hydrophobic regions, respectively. Polar water molecules and the protonation of nitrogen in the acridine ring (a) and side chain (b) are indicated as blue circles and red characters, respectively. Lower panel: The distribution (%) of these structures as a function pH are shown as blue (HP), green (CAD), and yellow (LP), respectively. The estimated pH-dependent octanol/water partition coefficient (logD) is indicated as a black line.

### QC alters to the structure inefficiently retaining photoenergy in acidic conditions

HP-QC and CAD-QC exist in the diprotonated (QC·2H^+^) and monoprotonated forms (QC·H^+^) by spectral analysis (*31*). However, it is unclear which structure the spectrum reflects and the significance of water on structural change. To determine these factors, we performed 3D fluorescence analysis (Fig. 2). QC had 3 major excitation peaks (270/360/410 nm) and one emission peak (500 nm) (Figs. 2A and S4). Because the excitation spectra are attributed to the number of conjugated double bonds (*32*), the acridine ring, consisting of 7 conjugated double bonds, is responsible for the spectrum of QC. The redshift of the emission wavelength is due to the Stokes shift, where QC loses excitation energy via radiationless deactivation by molecular motion (Fig. S3). The fluorescence intensities (Em 500) at the three excitation peaks increased with increasing pH (Fig. 2B), in accordance with the pH-dependent increase in the relative fluorescence quantum yield (*31*). Thus, deprotonation of nitrogen in the ring with a pH increase contributes to the fluorescence increase (Figs. 2B and S3A).

**Fig. 2.**
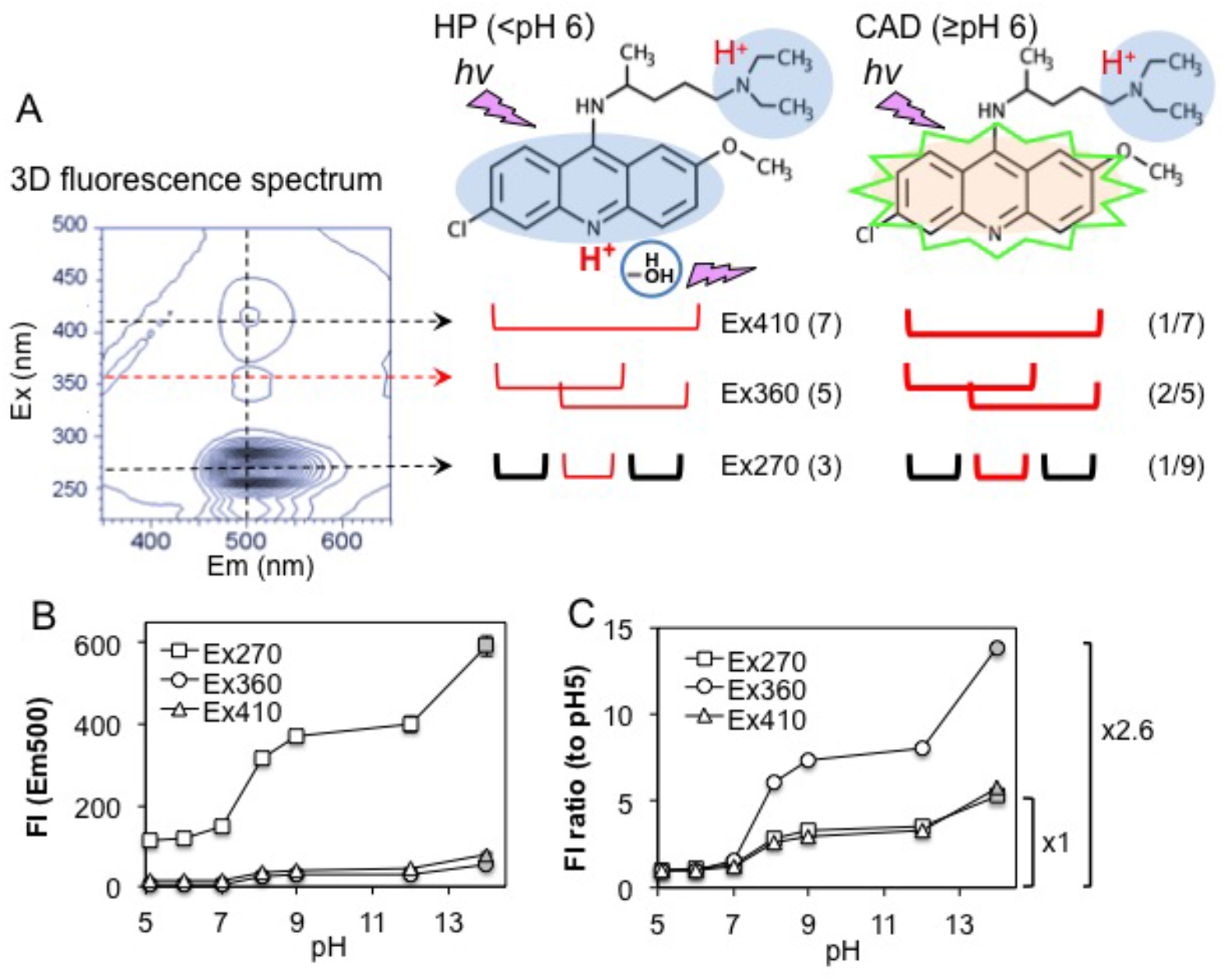
Relationship between the 3D fluorescence spectrum of QC and the structures. **(A)** Three-dimensional (3D) fluorescence spectra for the excitation (vertical line) and emission (horizontal line) of QC (5 μM) in 0.1 M phosphate buffer (pH 7.5) are shown in the left panel. The contour lines indicate the fluorescence intensity (FI). The HP and CAD forms of QC are shown on the right. The *hv* and green lines indicate the excitation energy and fluorescence, respectively. The structures absorbing energy around the indicated excitation peaks and the conjugated double bond numbers are indicated in the lower lines and parentheses (center). The ratios of protonated nitrogens per conjugated double bond number are indicated in parentheses (right). Red and black lines indicate the structures with and without protonated nitrogens, respectively. **(B)** The fluorescence intensities (FIs) of the emission (500 nm) at three excitation peaks are plotted against the indicated pH values (5.1, 6, 7, 8.1, 9, 12) and octanol (gray shapes). **(C)** The ratios of the FIs at each pH/pH 5 in B are plotted against the indicated pH values. Data are the mean (SE) (*n* = 3).

Apparent difference in the maximum emission wavelengths at Ex410 was not observed in water solutions with a pH decrease (Fig. S3B). Thus, the decrease in fluorescence with decreasing pH might not be typical radiationless deactivation. Presumably, the protonated nitrogen in the acridine ring would increase its electrical interaction with polar water molecules. If the interaction with water is responsible for the fluorescence decrease, the fluorescence should increase in nonpolar solutions. Indeed, the fluorescence at the three excitation peaks increased in an octanol solution (Fig. 2B), indicating that protonation of the acridine ring would partially dissipate the photoexcitation energy into the water as a quencher. Hence, QC would alter to the structure inefficiently retaining the photoenergy under acidic conditions.

### CAD-QC is formed at physiological pH

Three major excitation peaks (270/360/410 nm) reflect the number of conjugated double bonds in the acridine ring (*32*). Ex270 would excite the pyridine and two benzene structures (3 conjugated double bonds) in the ring (Fig. 2A). Because only the nitrogen of pyridine is protonated (1/3) but the two benzenes are not, Ex270 would reflect the effect of one protonation against the 9 conjugated double bonds (1/9) consisting of independent 3 structures. Ex360 would excite two redundant structures consisting of the quinoline structure (5 conjugated double bonds), Because each the nitrogen is protonated, Ex360 would reflect the effect of two protonation against the 5 conjugated double bonds (2/5). Ex410 would excite the whole acridine ring (7 conjugated double bonds) and reflect the protonation effect of 1/7. The acridine ring is almost completely protonated as HP-QC at pH 5, whereas it deprotonates with increasing pH, and is completely deprotonated as CAD-QC (LP-QC) in octanol, as estimated in Fig. 1B. Thus, the fluorescence ratios (Em 500) of each pH/pH 5 solution by each excitation peak would reflect the deprotonation/protonation (CAD/HP) ratio of the pyridine (Ex270), quinoline (Ex360), and acridine (Ex410) structures, and the ratios of octanol/pH 5 solution are the maximum CAD/HP ratios. These fluorescence ratios increased with increasing pH (Fig. 2C). In particular, the maximum CAD/HP ratio at Ex360 was 2.6-fold higher than those at Ex270 and Ex410. This is similar to the ratios of the protonation effect estimated by the structures Ex360/Ex270 (3.6:18/5) and Ex360/Ex410 (2.8:14/5). These results indicate that CAD-QC is formed at physiological pH and that Ex360 efficiently detects CAD-QC.

### HP-QC but not CAD-QC is excluded by the membrane system

To determine the pH effects on the interaction of QC with cells, the amount of QC adsorbing to yeast cells were determined at various pH values (5-9). In living cells, the cell-adsorbed fluorescence increased with the pH-dependent increase in CAD-QC (Fig. 3A), indicating that CAD-QC, but not HP-QC, is taken into cells. In cells treated with ethanol that destroyed the membrane structure or with formaldehyde that caused a loss in membrane protein function, the fluorescence increased at pH 5-7 but not at pH 8. This suggests that HP-QC is excluded by the membrane system at pH ≤ 7, and CAD-QC is taken into cells via the membrane system at pH 8. Because the fluorescence increased at pH 9 irrespective of treatment, LP-QC might also adsorb to insoluble materials such as the cell wall other than membranes.

**Fig. 3.**
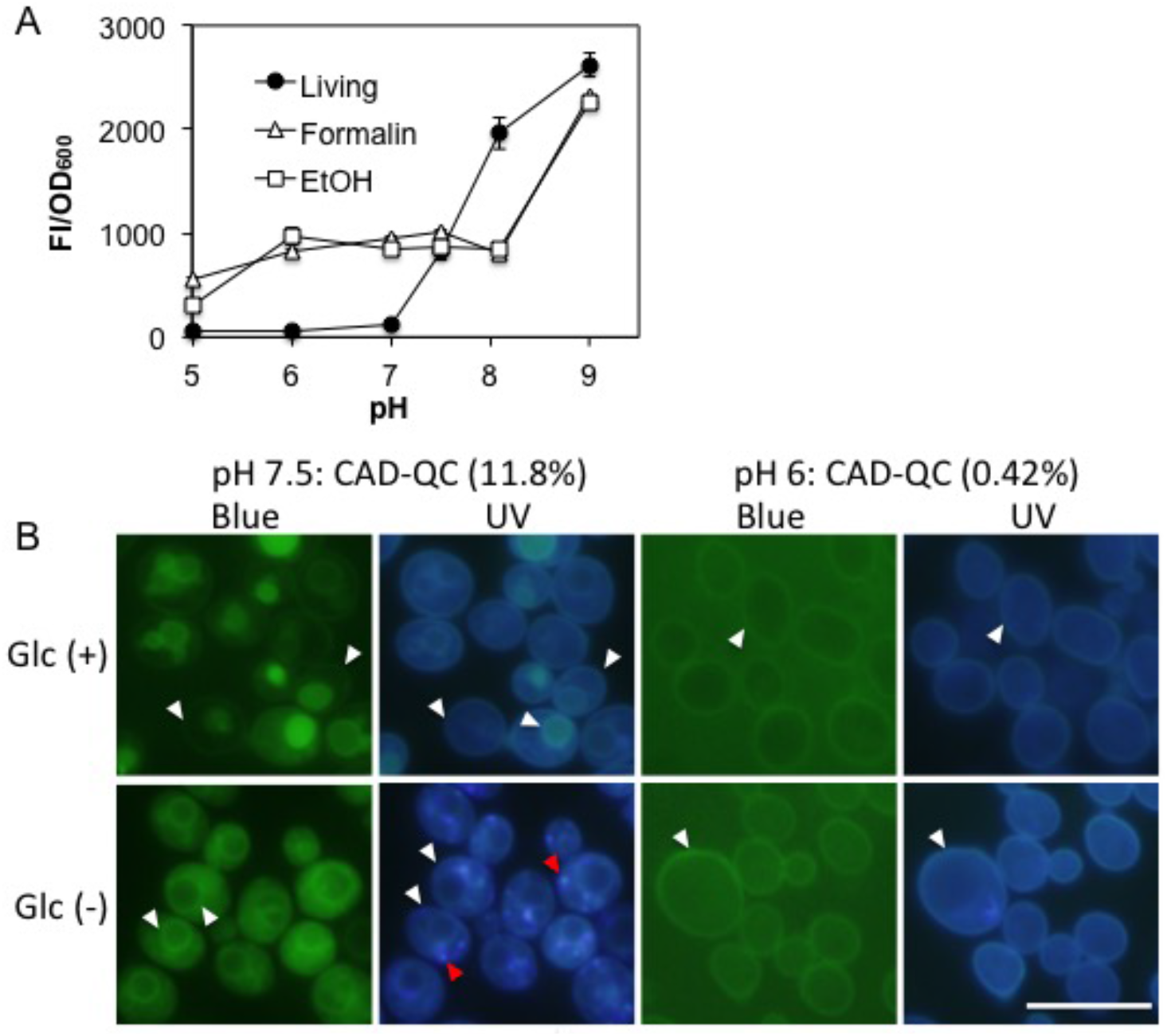
pH-dependent binding of QC with cells. **(A)** The QC fluorescence bound to BY4741 cells (FI/OD_600_) in buffer containing 2 % glucose at the indicated pH values was plotted against pH. Cells treated with or without formalin and ethanol are indicated. Data are the mean (SE) (*n* = 3). **(B)** Logarithmic BY4741 cells were suspended in phosphate buffer (0.1 M, pH 6 or 7.5) with or without 2% glucose and treated with 100 μM QC for 5 min. The estimated ratios of CAD-QC at each pH are indicated in parentheses. The fluorescence signals of QC were monitored by blue and UV irradiation, and the weak QC signals at pH 6 were detected by long exposure. The white arrows indicate the signals at membrane structures, and the red arrows indicate unknown granules formed by UV irradiation.

### CAD-QC localizes at membranes

We observed the localization of QC in yeast at pH 7.5 (CAD-QC: 11.8%) and pH 6 (CAD-QC: 0.4 %) (Fig. 3B). At pH 7.5 (2% glucose), the QC signal excited by 410 nm blue light was detected in acidic vacuoles and weakly in the plasma membrane. UV light at 360 nm preferentially detected signals in both plasma and vacuolar membranes, indicating that CAD-QC localizes in the membrane. At pH 6.0, weak QC signals were observed in the plasma membrane but not in vacuoles, irrespective of the excitation light and glucose, indicating that HP-QC is not efficiently taken into the cells as seen in Fig. 3A. At pH 7.5 without glucose required for vacuolar protonation, QC signals were observed in the plasma and vacuolar membranes, although unknown granules formed immediately upon UV exposure. Therefore, CAD-QC would penetrate into lipid membranes due to its amphiphilic structure.

### CAD-QC is crucial for antimicrobial activity

To determine the biological activity of QC, its minimum inhibitory concentration (MIC) was determined at the physiological pH (5-8) of yeast. The MIC is a relative value in response to the cell number. Under these conditions, using 1.25 × 10^5^ cells in 100 μl, the MICs were 50 mM (pH 6), 2.5 mM (pH 7), and 83 μM (pH 8). Yeast grew normally at 50 mM and pH 5 (CAD-QC: 0.04 %, HP-QC: 99.96 %), indicating that HP-QC does not have antimicrobial activity, as observed in the other microorganisms (*7*, *8*). This indicates that the antimicrobial activity of QC increased approximately 600-fold from pH 6 to 8 and 30-fold from pH 7 to 8 (Fig. 4A). Similarly, the antimicrobial activity of CQ increased approximately 30-fold from pH 7 (MIC: 15 mM) to pH 8 (MIC: 520 μM), although the yeast grew normally at 60 mM and pH 6 (Fig. S4). In contrast, because the MICs of CPZ without the HP structure were 250 μM (pH 6), 31 μM (pH 7), and 15 μM (pH 8), the antimicrobial activity of CPZ increased approximately 16-fold from pH 6 to 8 and 2-fold from pH 7 to 8. Thus, the drastic change in the antimicrobial activity of the antimalarial drugs is largely attributed to the pH-dependent structural change of CAD from HP, rather than biological mechanisms. Presumably, CAD-QC but not HP-QC exerts antimicrobial activity via nonspecific interactions with membranes.

**Fig. 4.**
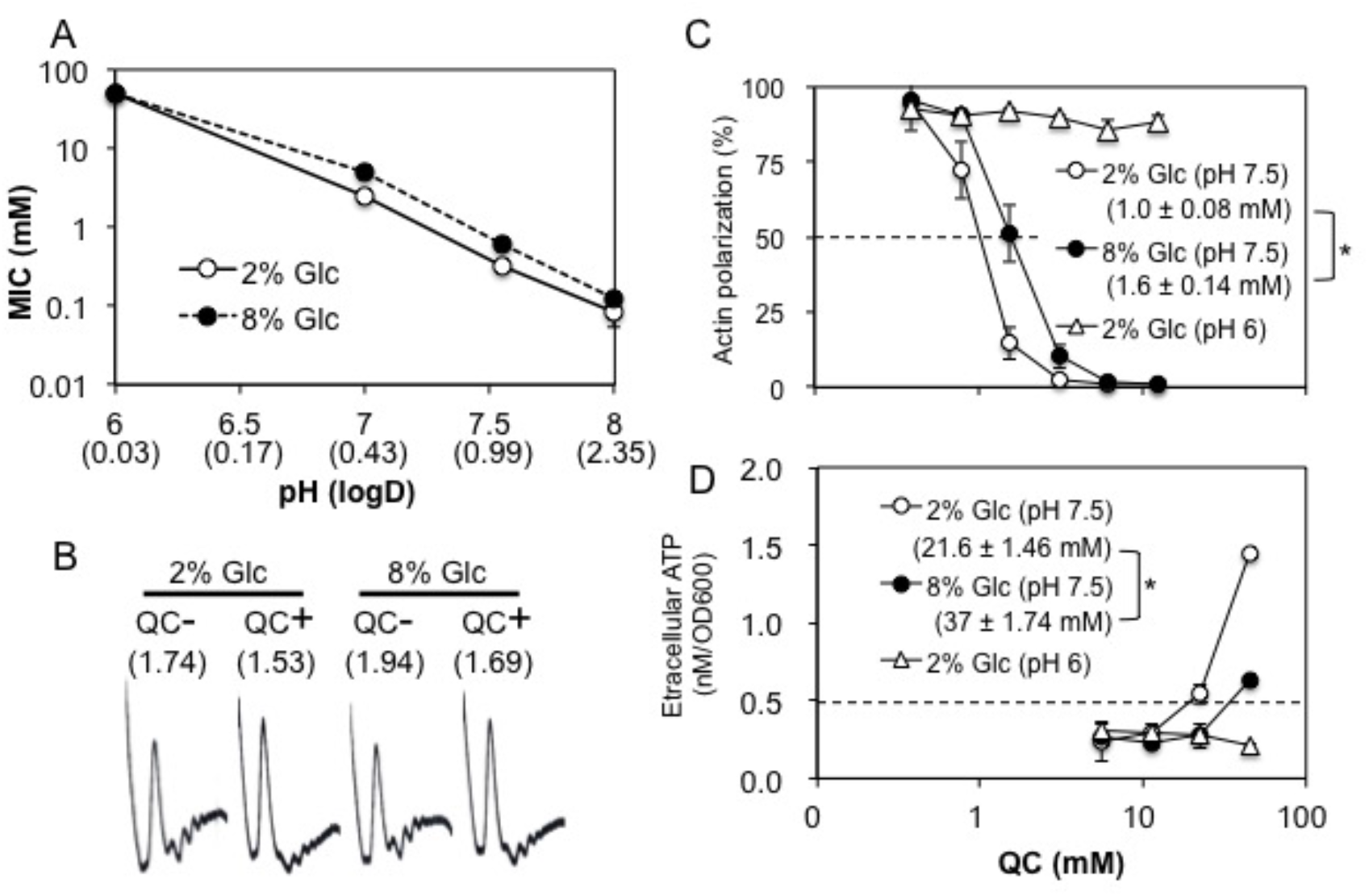
Biological activity induced by CAD-QC. **(A)** The MICs of QC for BY4741 cells were determined after 48 h in YPD containing the indicated amount of glucose buffered at the indicated pH. The estimated logD of QC at each pH is indicated in parentheses. Data are the mean (SE) (*n* ≥ 3). **(B)** Logarithmic BY4741 cells grown in YPD containing the indicated glucose concentrations (pH 7.5) were treated with 1.5 mM QC for 30 min. The polysome/monosome (P/M) ratios are indicated in parentheses. Representative data from one of three independent experiments are shown. **(C)** Logarithmic BY4741 cells were incubated for 1 h in YPD containing the indicated glucose concentrations buffered at the indicated pH values and then treated with QC for 30 min. The percentage of cells with polarized actin in the total counted cells are plotted against the QC concentration. Half inhibitory concentrations of actin polarization are indicated in parentheses. Data are the mean (SE) (*n* = 3), *\*p* = 0.02. **(D)** ATP amounts in the supernatants of the cultures treated in panel C are plotted against the concentration. The concentrations of QC where the ATP amount was 0.5 nM/OD_600_ are indicated in parentheses. Data are the mean (SE) (*n* = 3), *\*p* = 0.015.

### CAD-QC induces glucose-starved reactions via membrane interactions

Excessive amounts of glucose alleviate the antimicrobial activities of yeast induced by CADs, such as the local anesthetic tetracaine and CPZ, irrespective of structural differences (*26*). Thus, it might also alleviate the effects induced by CAD-QC. The MICs at 8% glucose were 2-fold higher than those at 2% glucose at pH 7-8 (CAD-QC: 4-29.6 %), whereas no change in MIC was observed by excessive glucose at pH 6 (CAD-QC: 0.4 %) (Fig. 4A). This suggests that CAD-QC induces glucose starvation. The above CADs inhibit translation initiation and actin polarization in yeast as glucose-starved reactions and lyse the cells at high concentrations via amphiphilic structures (*26*, *33*). QC inhibited translation initiation at pH 7.5, which was slightly alleviated by 8% glucose, as judged by polysome/monosome ratios (Fig. 4B). QC also inhibited actin polarization in a dose-dependent manner at pH 7.5 but not pH 6.0, and this inhibition was partially alleviated by 8% glucose, as judged by the half inhibitory concentration(IC_50_)(Fig. 4C). Similarly, QC lysed yeast cells at high concentrations and was slightly alleviated by 8% glucose, as judged by extracellular ATP amounts (Fig. 4D). Thus, CAD-QC but not HP-QC induced glucose-starved reactions in yeast via membrane interactions, and eventually destroyed the membrane at high concentrations.

### CAD-QC noncompetitively inhibits hexose transport

Because hexose transporters localize at plasma membranes to take up environmental hexoses (*26*, *34*, *35*), CAD-QC might inhibit transporter function as a membrane target. We evaluated the effect of QC on the initial rate of 2-deoxyglucose (2DG) uptake (zero-trans influx) in the glucose-starved HXT2m strain expressing only a high-affinity hexose transporter. QC inhibited 2DG uptake in a dose-dependent manner at pH 7.5 more efficiently than at pH 6, as judged by the IC_50_ (Fig. 5A). Thus, CAD-QC strongly inhibits the function of hexose transporters on the plasma membrane. To determine the inhibition mechanism of Hxt by CAD-QC, the kinetic parameters for 2DG uptake inhibition were determined at pH 7.5 (Fig. 5B). In the absence of QC, the Km value of Hxt2 was approximately 0.8 mM, which was similar but slightly less than the value (1.5 mM) determined by the radioisotope method in the *hxt1-7* null mutant expressing the *HXT2* gene (*36*). The Vmax value (710 nmol/min·mg cells, equivalent to 208 nmol/min·OD_600_) was higher than that in a previous report (97 nmol/min·mg cells) (*36*), possibly because of the high expression of *HXT2* on the multicopy vector. The Vmax values decreased with increasing QC, but the Km values did not, indicating that QC inhibits 2DG transport of Hxt2 noncompetitively at pH 7.5. Therefore, CAD-QC would inhibit Hxt2 function by interacting with sites other than the hexose recognition site.

**Fig. 5.**
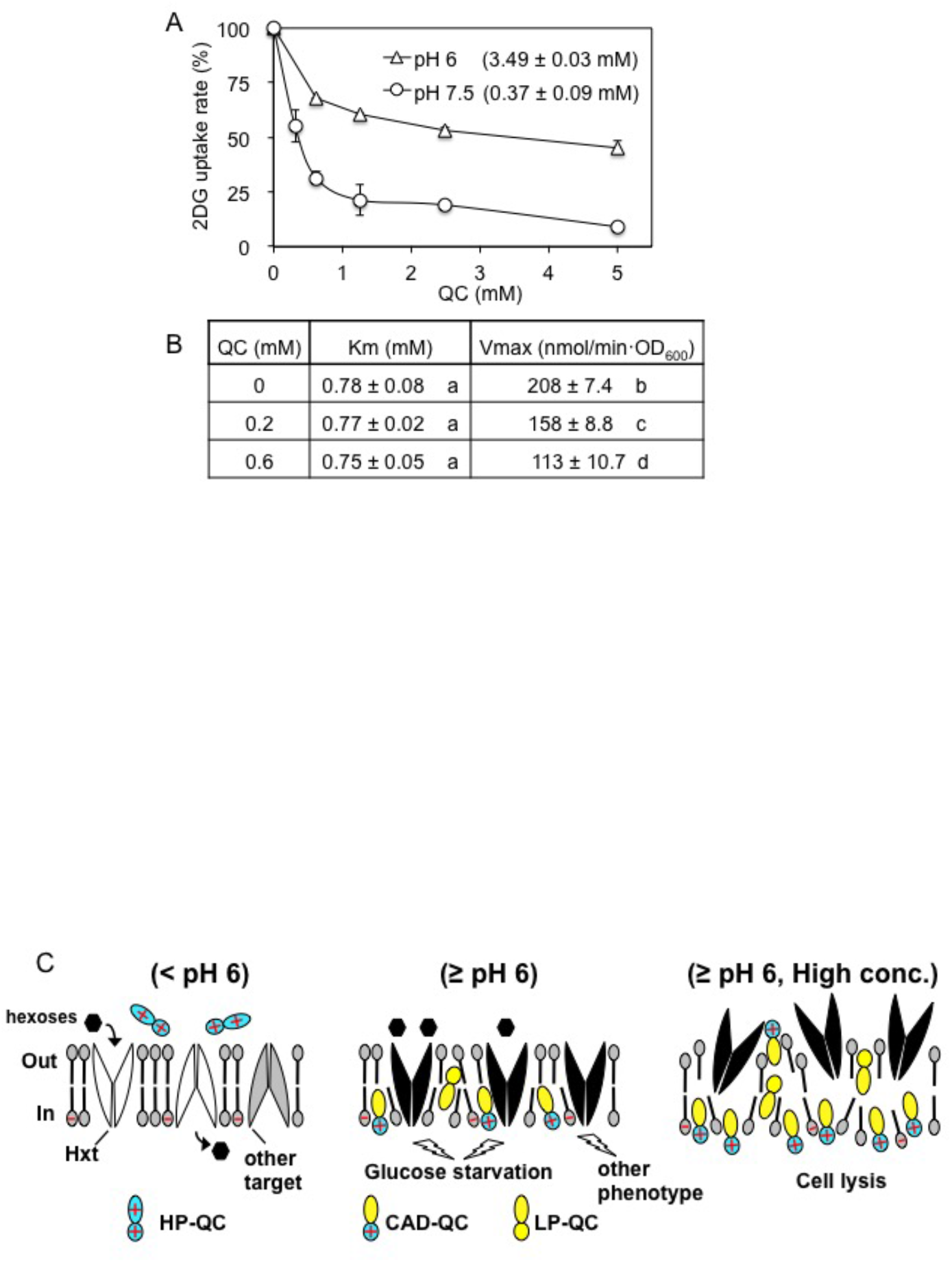
The inhibition mechanism of Hxt via nonspecific interaction of QC with membrane. **(A)** Inhibition of 2DG uptake by QC. The initial rates of 2DG uptake at each QC concentration were plotted as the ratio to the rate without QC. The half inhibitory concentrations (IC_50_) in each buffer are indicated in parentheses. **(B)** Kinetic parameters for 2DG uptake inhibition in the HXT2m strain. Data are the mean (SE) (*n* ≥3). Different letters denote significant differences, *p* <0.05. **(C)** Models for Hxt inhibition caused by nonspecific interaction of QC with membranes. (Left) HP-QC does not have antimicrobial activity. Active Hxts and the other target are indicated as white and gray shapes, respectively. (Center) CAD-QC adsorbs to the cells and then stably associates with the negatively charged inner leaflets of the lipid bilayer by converting to LP-QC at the center of the bilayer. Nonspecific interactions of CAD-QC with the membrane would affect various membrane targets (black shapes) and noncompetitively inhibit the function at the lateral side of abundant Hxts with high probability, resulting in preferential glucose starvation. (Right) Increasing drug dose destroys the membranes by excessively penetrating into the lipid layer.

## Discussion

Various action and resistance mechanisms of antimalarial drugs have been studied regarding the biological aspects of parasites (*2*, *37*). However, we focused on pH-dependent structural changes in the drug mechanisms and categorized the structures of QC into the HP, CAD, and LP forms. Basically, the interaction of QC with cells obeyed the pH partition hypothesis: ionized drugs diffuse through biological membranes primarily in their unionized form, and the extent of drug absorbed across membranes depends on the degree of ionization (*38*). However, this hypothesis considers the action of the ionized drug in the cytoplasm but not in the membrane. Although structural changes in ionized QCs are known as QC·2H+ and QC·H+ (*31*), their biological significance remains unknown. In addition, it seems that the obvious accumulation of these ionized QCs into acidic organelles confused the elucidation of the action mechanism. Here, we describe the importance of the CAD form on biological activity.

### Nonspecific interaction of CAD-QC with the membrane inhibits various membrane targets, including hexose transporters

HP-QC easily dissolves in water but hardly interacts with insoluble materials such as the cell wall and membranes (Figs. 1B and 3A); thus, it would hardly have antimicrobial activity (Fig. 4A). CAD-QC could easily adsorb to cells (Fig. 3A) and penetrate nonspecifically into membrane lipids via the CAD structure (Fig. 3B). It would stably interact with the negatively charged inner leaflets of the lipid bilayer via electrostatic interactions, as CPZ does (*26*, *28*, *29*), and affect various membrane targets, thereby exerting strong antimicrobial activity (Fig. 4A). Since the HP and CAD forms exist as mixtures at the physiological pH values (5-8) of yeast, the antimicrobial activity is determined by the pH-dependent abundance ratio of CAD-QC (Fig. 1B). Although LP-QC does not exist at pH 5-8 (Fig. 1B), CAD-QC would convert into LP-QC at the waterless center of the lipid bilayer during translocation into the cytoplasm. Thus, LP-QC would also exist in the cell membrane in proportion to the extracellular CAD-QC abundance. Consequently, the interaction of the CAD form with the membrane is crucial for its pharmacological effects such as antimicrobial activity.

The nonspecific interaction of QC with the membrane affects numerous membrane targets, thereby exhibiting various phenotypes, as observed for various CADs (*4*, *39*). Nevertheless, the hexose transporter is an important membrane target for QC in yeast because additional glucose alleviates the antimicrobial activity (Fig. 4A) and QC induces glucose starvation reactions (Fig. 4B and C) and inhibits the uptake of 2DG (Fig. 5). Because the yeast hexose transporter is the largest family encoded by 18 genes, Hxts would be abundant in the plasma membrane (*34*). Thus, the nonspecific interaction of QC would noncompetitively inhibit the function at the lateral side of abundant Hxts with high probability (Fig. 5), resulting in preferential glucose starvation according to the lipid theory (*26*). Although QC is eventually translocated into acidic vacuoles, inhibition would be maintained as long as QC exists outside the cells. Practically, additional glucose increases the MICs at pH 7-8 only two-fold (Fig. 4A). Additionally, the IC_50_ of QC for Hxt2 inhibition at pH 8 was approximately 1/10 of that at pH 6 (Fig. 5A), although the MIC of QC at pH 8 was approximately 1/600 of that at pH 6 (Fig. 4A). These results indicate that CAD-QC affects antimicrobial targets other than Hxt. Because CQ and QN, having the CAD structure (Fig. S5), are known to inhibit the transport of iron (*10*), tryptophan, tyrosine (*11*), glucose (*12*), and thiamine (*13*) in budding yeast, CAD-QC might also inhibit the transport of these nutrients. Furthermore, the local anesthetic tetracaine, a CAD, increases the membrane targets with the dose increase and finally destroys the membrane (*40*), similar to CAD-QC (Fig. 4D). Since these effects would result from nonspecific interactions of the CAD structure with the membrane, the mode of inhibition of other targets might be similar to the inhibition mechanism of Hxt2. These ideas are summarized in Fig. 5C.

This mechanism would be applicable for various organisms having membranes and exhibit two pharmacological effects. First, nonspecific interactions with the membrane would inhibit the functions of various membrane proteins, including the nutrient uptake system, thereby inducing nutrient starvation. This would suppress cellular function and might be responsible for the pleiotropic pharmacological effects, such as antimalarial, antifungal, anticancer, and immunosuppressive effects (*2*–*15*). Second, membranes containing foreign compounds such as QC and CQ do not have normal functions for membrane fusion/separation between the host and parasites/enveloped viruses, thereby exhibiting antimalarial and antiviral activities (*2*, *4*–*6*). Because this mechanism is also based on nonspecific interactions of the CAD structure with the membrane, CADs might be effective drugs to suppress parasitic/microbial/viral infections irrespective of protein alteration by gene mutation.

### A decrease in the CAD structure in acidic environments accounts for drug resistance

The Meyer-Overton correlation was discovered over 120 years ago and states that the anesthetic potency of various anesthetics increases logarithmically with lipophilicity (*41*). This is a prototype for the quantitative structure-activity relationship (QSAR) in which the biological potency of various drugs increases logarithmically with lipophilicity/hydrophobicity (*42*), although it remains unclear whether drugs target lipids or proteins. The MICs of QC and CQ decreased logarithmically with logD increases from pH 5-8 (Fig. S5). This result indicates that the same compound with various lipophilicities obeys the Meyer-Overton correlation, thereby avoiding complicated interpretations due to various structures. To put the correlation differently, the drastic logarithmic decrease in MIC would be determined by the amount of the CAD structure interacting with membrane lipids, thereby supporting lipid theory.

The MICs of QC, CQ and CPZ were determined from linear QSAR models regressed as a function of pH (Fig. S5B). According to these models, the dose ratios equivalent to the MICs at pH 7.4 were plotted around the normal human blood pH (Fig. 6A). Similar ratio changes were observed for QC and CQ, but not CPZ without the HP structure. This indicates that a slight change in pH at the clinical level would largely affect the effects of the antimalarial drugs. For example, for the effects equivalent to pH 7.4, pH 7.2 and pH 7.6 would require 2-fold and one-half the dose, respectively. Presumably, the ratios would be similar to those for the inhibition of parasitic/viral infections because these ratios are attributed to the pH-dependent structural change of CAD from HP, although precise estimation of the ratio might require a model between pH and these phenomena. Hence, these antimalarial drugs would show a reduction in their pharmacological effects with a slight decrease in blood pH. Various quinoline antimalarial drugs would also be pH-dependent CADs because their nitrogen atoms are protonated (Fig. S5) (*2*). Computational analysis estimated the pH distributions of their CAD structures (Fig. 6B). According to the distributions, because the CAD structure of HCQ decreased with a slight pH decrease, the pharmacological effects are also predicted to decrease, as in the case of QC and CQ, although the other quinoline drugs showed different distribution patterns.

**Fig. 6.**
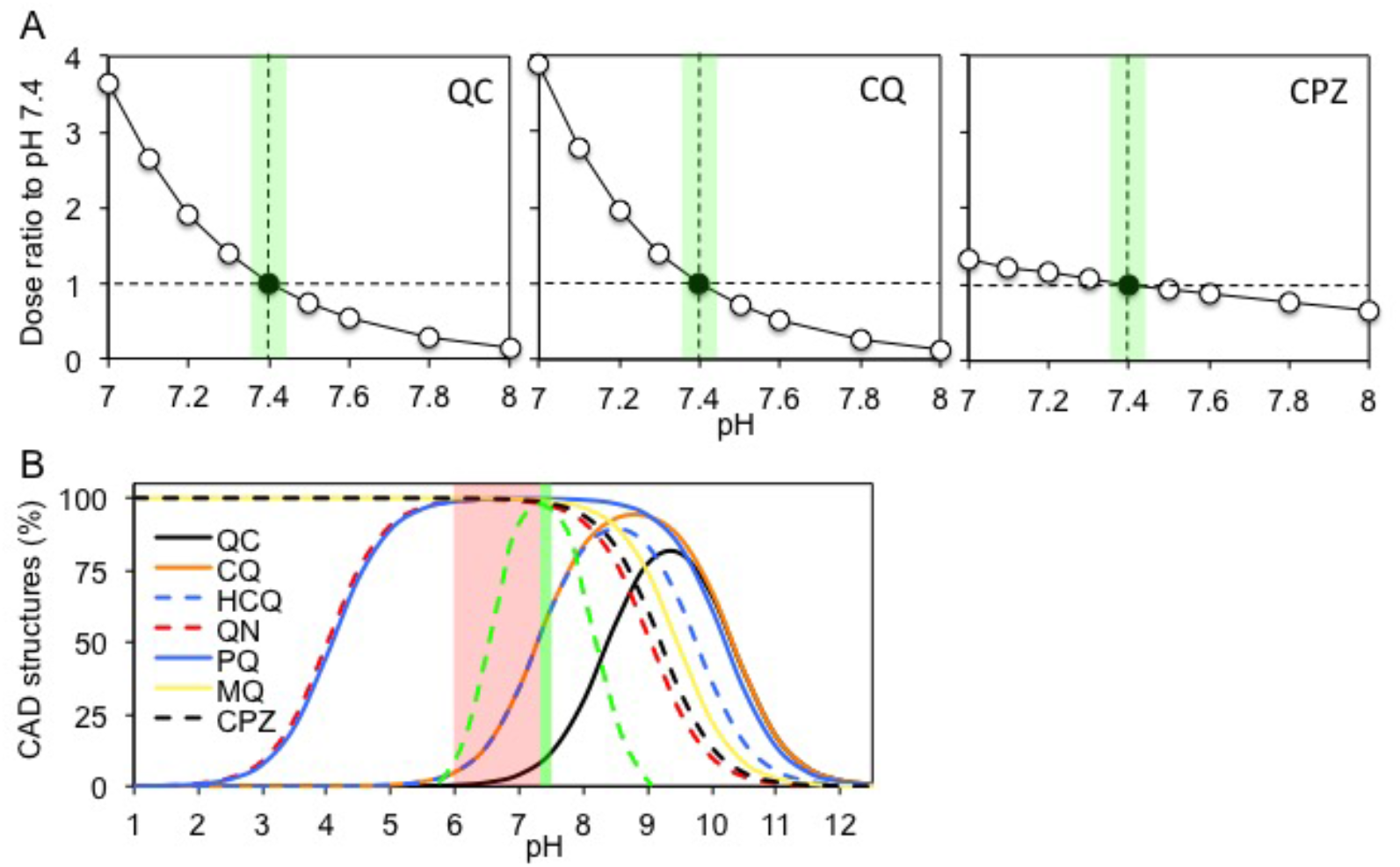
The activities of antimalarial drugs change with pH. **(A)** The dose ratios equivalent to the MICs of QC, CQ, and CPZ at normal human blood pH (7.4) were plotted at the range of pH 7-8, according to the linear models in Fig. S4. **(B)** The distributions of the CAD structures of various antimalarial drugs and the antipsychotic CPZ at different pH values were estimated using MarvinSketch. Green and red zones indicate normal and abnormal acidic ranges of human blood pH, respectively. The green dotted line indicates the CAD distribution of the ideal drug to treat for blood infection. Abbreviations, QC: quinacrine, CQ: quinacrine, HCQ: hydroxychloroquine, QN: quinine, PQ: primaquine, MQ: mefloquine, CPZ: chlorpromazine.

There is a variety of pH values in human tissues, such as gastric acid (< 3.5), urine (6), the liver (7.2), blood (7.4), pancreatic secretions (8.1), and the brain (7) (*43*, *44*). Thus, the potency of quinoline drugs evaluated at a simple pH would be different from those in the whole body with complicated pH values. Additionally, microorganisms such as yeast and cancer cells produce acidic metabolites, thereby lowering the pH of their microenvironments (*7*, *8*, *45*). In these acidic environments, several quinoline drugs would not act efficiently. For example, falciparum malaria is known to be resistant to CQ. Because the common manifestation of falciparum malaria is acidosis irrespective of age (*3*), CQ would not act efficiently on falciparum malaria causing acidic blood. Hypnozoites of vivax and ovale malaria in the liver are also resistant to CQ (*3*). This is also possibly due to the low pH (7.2) of the liver. Importantly, although these cases are recognized as drug resistance, there would be a decrease in drug potency from structural alterations in acidic environments. This should be distinguished from biological mechanisms such as drug efflux. Interestingly, quinine (QN) and primaquine (PQ) are effective for falciparum malaria and hypnozoites, respectively (*3*). Because these drugs form the CAD structure over a wide pH range (Fig. 6B), they would act effectively even in acidic environments. Mefloquine (MQ) is also effective for falciparum malaria but has adverse psychiatric effects (*3*). Because the pH distribution of the CAD structure is similar to that of the antipsychotic CPZ (Fig. 6B), MQ acts on the brain with a low pH similar to CPZ. This suggests that drugs with a CAD structure over a wide pH range induce unexpected adverse effects by persistently interacting in tissues with various pH values. Therefore, it is important to know the pH at the pathogenic site and find/design a drug forming the CAD structure only at the pH of the pathogenic site. Presumably, this would create a useful drug with few adverse effects.

### The way to use antimalarial drugs efficiently

Administration doses of the drugs would be relatively determined by their lipophilicity (logP, logD) according to the Meyer-Overton correlation. However, it is important to take into account pH when using pH-dependent CADs, including antimalarial drugs. When the pathogenic site is the blood and the patient’s blood pH is higher than 7.4 from alkalosis, the dose of antimalarial drugs such as QC, CQ and HCQ should be reduced because of increased potency and toxicity (Fig. 4C and D). When the pH is lower than 7.4, such as in falciparum malaria (*3*) and COVID-19 pneumonia patients with acidosis, the conventional dose would not reach the minimum effective concentration. In these cases, the drug dose should be increased. However, overdose might cause adverse effects by adsorption into alkaline tissues other than vessels (Fig. 3A). To avoid this, achieving normal blood pH by clinical treatment might be effective at the conventional dose of antimalarial drugs. Alternatively, QN, PQ, and MQ with a wide pH range of the CAD structure might be effective irrespective of acidosis (Fig. 6B), although attention to their adverse effects is required.

## Acknowledgments

We thank Drs. Y. Inoue, W. Nomura, and K. Yamada of Kyoto University; T. Kasahara of Teikyo University; E. Boles of Goethe University; and Y. Osakabe of Tokushima University for providing plasmids and strains. Supporting data are available in the supplementary materials.

## Supplementary Materials

Materials and Methods

Figs. S1 to S5

References (*46*–*52*)

