## Supplemental Materials for "Antimalarial drugs lose their activity with a slight drop in pH"

**This PDF file includes:**

Materials and Methods

Figs. S1 to S5

References (*46-52*)

### Materials and Methods

#### Strains, media, plasmids, and growth conditions

Standard genetic manipulation of yeast and DNA manipulation were performed as described previously (*46*). The media used for the budding yeast *Saccharomyces cerevisiae* were YP (1% yeast extract and 2% polypeptone) and YPD (YP containing 2% glucose). The yeast strain used was BY4741 as the wild-type (MATa *his3Δ1 leu2Δ0 ura3Δ0 met15Δ0*) and EBY.S7, deleting all 18 hexose transporter genes (MATa *hxt1-17Δ gal2Δ agt1Δ stl1Δ leu2-3, 112 ura3-52 trp1-289 his3-Δ1 MAL2-8c SUC2 fgy1-1*). EBY.S7 harboring Hxt2mnx-pVT, the *ADH1* promoter–driven *HXT2* gene on multicopy vector pVT102-U (*2µ ori*, *URA3*), was used as the HXT2m strain expressing a high-affinity hexose transporter alone (*47, 48*). Cellular turbidity ($OD_{600}$) was monitored by a UV spectrophotometer (Ultrospec 1100 Pro, GE Healthcare, USA), and the cell numbers were counted with hematocytometer. The values were determined in YPD cultures of BY4741 ($OD_{600}$ = 1:2.5 × $10^7$ cells/ml) and HXT2m ($OD_{600}$ = 1:1.2 × $10^7$ cells/ml).

#### pH adjustment of solution

One molar solutions of sodium acetate, sodium phosphate, Tris-HCl, and HEPES-NaOH buffers were prepared to obtain the final pH values (5.1, 6, 6.5, 7, 7.5, 8.1, 12, and 9) at 0.1 M. Because final pH values at 0.1 M vary in the solutions for use, the 1 M solutions were prepared for water and YPD solutions, respectively. To avoid a decrease in pH by glycation of YP medium containing 8 % glucose, YP and glucose were separately autoclaved and then mixed with 1 M sodium phosphate buffer to obtain the final pH values (6 and 7.5) at 0.1 M.

**Chemicals**

Quinacrine (QC) and the antipsychotic chlorpromazine (CPZ) were purchased from Sigma-Aldrich (St. Louis, MO, USA). Chloroquine (CQ), 2-deoxy-D-glucose (2DG), and cycloheximide (CYH) were purchased from Wako Pure Chemicals (Tokyo, Japan). Rhodamine phalloidin was purchased from Molecular Probes (Eugene, OR, USA).

**Computational methods**

The acid dissociation constants (pKa values) and distribution coefficients (logD values) were estimated using the MarvinSketch software package (19.8.0) (ChemAxon, Hungary) (http://www.chemaxon.com). The consensus logP method was applied to calculate the distribution coefficient of each molecule.

**Fluorescence analysis**

Fluorescence spectra for the excitation and emission of quinacrine (QC) were measured with a fluorescence spectrophotometer (model F-2700, Hitachi, Tokyo, Japan) in the indicated buffer solutions and octanol solution, respectively. The photomultiplier voltages were 250 V for the three-dimensional (3D) fluorescence spectra and 400 V for the cell-binding assay. The slit widths for excitation and emission were 10 nm.

**Cell binding assay**

Logarithmic cultures of the BY4741 strain in YPD were washed twice with water and suspended in buffers containing 2 % glucose with 0.1 M sodium acetate (pH 5.1), sodium phosphate (pH 6, 7, 7.5, and 8.1), or Tris-HCl (pH 9). Formaldehyde was added directly into the YPD culture at a final concentration of 5 % and incubated for 30 min. Ethanol (70%) was added into the cell pellets after the removal of YPD medium and mixed for 1 min. These chemically fixed cells were washed as above. The cell suspensions were treated as 100 μl of reaction mixture containing 2.5 × $10^7$ cells and 1 mM QC and incubated for 15 min at 25°C. The QC-treated cells were washed twice with the same pH buffer and then suspended in 1 ml of sodium phosphate buffer (0.1 M, pH 7.5). The fluorescence and cellular turbidity ($OD_{600}$) were measured using a fluorescence spectrophotometer and UV spectrophotometer, respectively.

**Determination of the minimum inhibitory concentration**

The minimum inhibitory concentration (MIC) of yeast was determined as described previously (*49*). Logarithmic BY4741 cells were mixed with YPD (0.1 M: pH 5.1, 6, 7, 7.5, or 8.1) containing 2 % or 8 % glucose at a final $OD_{600}$ of 0.05, and then 100 μl (1.25 × $10^5$ cells) was dispensed into 96-well microtiter plates (Corning, NY, USA). QC was added directly to the cell suspensions, 2-fold serial dilutions were performed in the wells, and the plates were incubated at 25°C for 48 h. The MIC was the lowest concentration that completely inhibited microbial growth, as judged by visual observation.

**Microscopic analysis**

For the detection of CPZ and QC, logarithmic BY4741 cells grown in YPD were washed with sodium phosphate buffer (0.1 M, pH 6 or 7.5) with or without 2 % glucose, suspended in the same buffer solution and then treated with drug at the indicated concentrations and times. Actin staining and calculation of the percentages of polarization were performed as described previously (*50*). Logarithmic BY4741 cells were incubated at 25°C for 1 h in YPD (0.1 M, pH 6 or 7.5) containing 2 % or 8 % glucose. Two hundred microliters of these cultures ($OD_{600}$ = 2.0) were treated with the indicated concentrations of QC at 25°C for 30 min. The cultures were fixed by adding formaldehyde at a final concentration of 5 % for 30 min, washed twice with phosphate-buffered saline (pH 7.5), and then stained with rhodamine phalloidin. The percent polarization was calculated as the fraction of cells exhibiting a polarized actin cytoskeleton in small- and medium-budded cells (n >300). Cells with <50% of their actin patches in the bud were considered to have a depolarized actin cytoskeleton. Images were recorded using a BX50 microscope with a DP71 CCD camera using DP controller software (Olympus, Tokyo, Japan) (*5*). Olympus U-MWU, U-MWIB/GFP, and U-MWIG cubes were used to detect CPZ, QC, and actin, respectively.

**Cell lysis assay**

Cell lysis was evaluated by the extracellular ATP amount leaked into the supernatant of the drug-treated culture. One hundred microliters of these cultures ($OD_{600}$ = 2.0) were treated with the indicated concentrations of QC at 25°C for 30 min. The reaction mixtures were centrifuged, and 10 μl of each supernatant was diluted with 90 μl of sodium phosphate buffer (0.1 M, pH 7.5). Five microliters of each sample was mixed with equivalent amounts of the luciferase/luciferin mixture from the Kinshiro ATP extraction system kit (LL-100-1; Toyo Ink, Tokyo, Japan), and ATP amounts were measured using a luminometer (AB-120, ATTO, Tokyo, Japan) according to the manufacturer's instructions. The extracellular ATP levels were calculated as [ATP concentration]/[$OD_{600}$].

**Polysome analysis**

Polysome analysis was carried out as described previously (*50*).

**Determination of the uptake rate of 2DG**

The initial rate of hexose uptake was determined under zero-trans entry conditions with 2-deoxyglucose (2DG) as substrate using the bioluminescent assay, which is based on measuring 2DG6P converted from 2DG transported via a hexose transporter (Glucose uptake-Glo$^{TM}$ assay, Promega). The manufacturer's method was improved for budding yeast as follows. Logarithmic cultures of the HXT2m strain grown in YPD were washed twice with sodium phosphate buffer (0.1 M, pH 7.4 and pH 6.0) and suspended in the same buffer as the glucose-starved cells ($OD_{600}$ = 2). Ten microliters of QC was mixed sequentially with 80 µl of the cell suspension at the indicated concentrations and incubated correctly for 10 min at 25°C. Then, 10 µl of 2DG was sequentially mixed with the QC-treated cells at the indicated concentrations (100 µl total, $2.4 \times 10^6$ cells) and incubated for 10 seconds at 25°C. The reaction mixtures were snap-frozen in liquid nitrogen. The dissolved mixtures were centrifuged at 4°C, and the cell pellets were mixed with 44 µl of 10 % $HClO_4$ for 10 min on ice. The acid extracts containing cells were neutralized with 6 µl of 5 M $K_2CO_3$ for 10 min on ice. After centrifugation, the supernatants of the neutralized solutions containing 2DG6P were diluted 20-fold in sodium phosphate buffer (0.1 M, pH 7.4), and the salt pellets were dissolved in 1 ml of water to measure the $OD_{600}$. Five microliters of the diluted solution was mixed with equivalent amounts of the 2DG6P detection reagent and incubated at 25°C for 30 min, and then the amounts of 2DG6P were determined using a luminometer (AB-120) according to the manufacturer's instructions. The amount of 2DG6P arising from the transported 2DG was normalized by subtracting the background luminescence arising from authentic G6P of the yeast cells untreated with 2DG. The initial rate of 2DG uptake was expressed as the unit per cell (nmol/min·$OD_{600}$). For the half inhibitory concentrations ($IC_{50}$), the initial rates were determined at the QC concentration (0-5 mM) in the presence of 1 mM 2DG. For kinetic parameters, the initial rates were determined in the presence of 2DG (0–4 mM) and QC (0–0.6 mM). It should be noted that this method was not applicable for the low affinity hexose transporter, possibly because a high concentration of transported 2DG inhibited hexokinase activity.

**Data analysis**

The $IC_{50}$, Km, and Vmax values were determined by the nonlinear least-squares method of the log-logistic (*51*) and Michaelis-Menten (*52*) models using the Solver add-in bundled with Microsoft Excel. The QC concentrations where the ATP amount was 0.5 nM/$OD_{600}$ were determined according to the linear models between two points including the value. Statistical analyses were performed by unpaired Student's *t*-test and one-way analysis of variance with the Tukey-Kramer multiple comparison test, using the R statistical software package (ver. 3.5.1; R Development Core Team 2018).

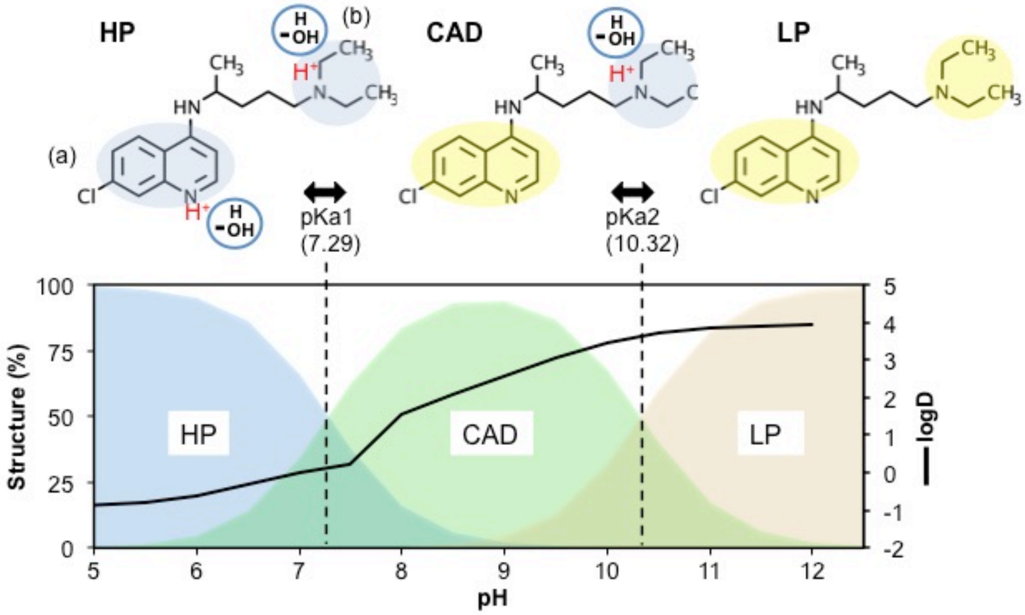


**Fig. S1. pH-dependent change of the structure and physicochemical properties of CQ**

(Upper) Changes in the structure and physicochemical properties as a function of pH were estimated using MarvinSketch. The pH-dependent CQ structures are shown as the hydrophilic (HP), cationic amphiphilic (CAD), and lipophilic (LP) forms. The blue and yellow areas are the same as those in Fig. 1B. Polar water molecules and protonation of the nitrogen in the quinoline ring (a) and side chain (b) are indicated as blue circles and red characters, respectively. (Lower) The distribution (%) of these structures with pH are shown as blue (HP), green (CAD), and yellow (LP). The estimated pH-dependent octanol/water partition coefficient (logD) is indicated as a black line.

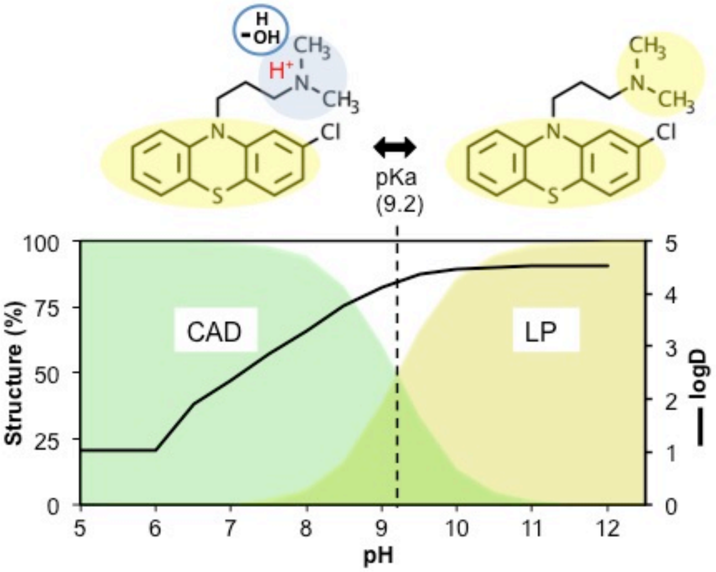


Fig. S2

**Fig. S2. pH-dependent change of the structure and physicochemical properties of CPZ**

(Upper) Changes in the structure and physicochemical properties with pH were estimated using MarvinSketch. The pH-dependent CPZ structures are shown as the cationic amphiphilic (CAD) and lipophilic (LP) forms. Blue and yellow areas are the same as those in Fig. 1B. Polar water molecules and protonation of the nitrogen on the side chain are indicated as blue circles and red characters, respectively. (Lower) The distribution (%) of these structures with pH are shown as green (CAD) and yellow (LP). The estimated pH-dependent octanol/water partition coefficient (logD) is indicated as a black line.

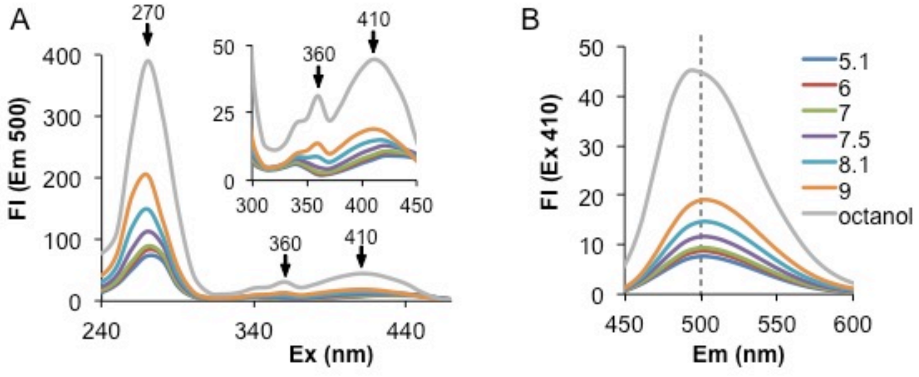


**Fig. S3. Spectral changes of the QC fluorescence with changes in pH**

Excitation spectra at the emission peak of 500 nm **(A)** and emission spectra at the excitation peak of 410 nm **(B)** of QC (5 μM) were measured in the indicated pH (5.1, 6, 7, 7.5, 8.1, and 9) solutions (0.1 M) and octanol. The expanded excitation spectra are shown in the upper panel in A. The arrows in A indicate the three excitation peaks in Fig. 2A. The dotted line in B indicates the emission peak at 500 nm. The photomultiplier voltage was 250 V, and the slit widths for excitation and emission were 10 nm.

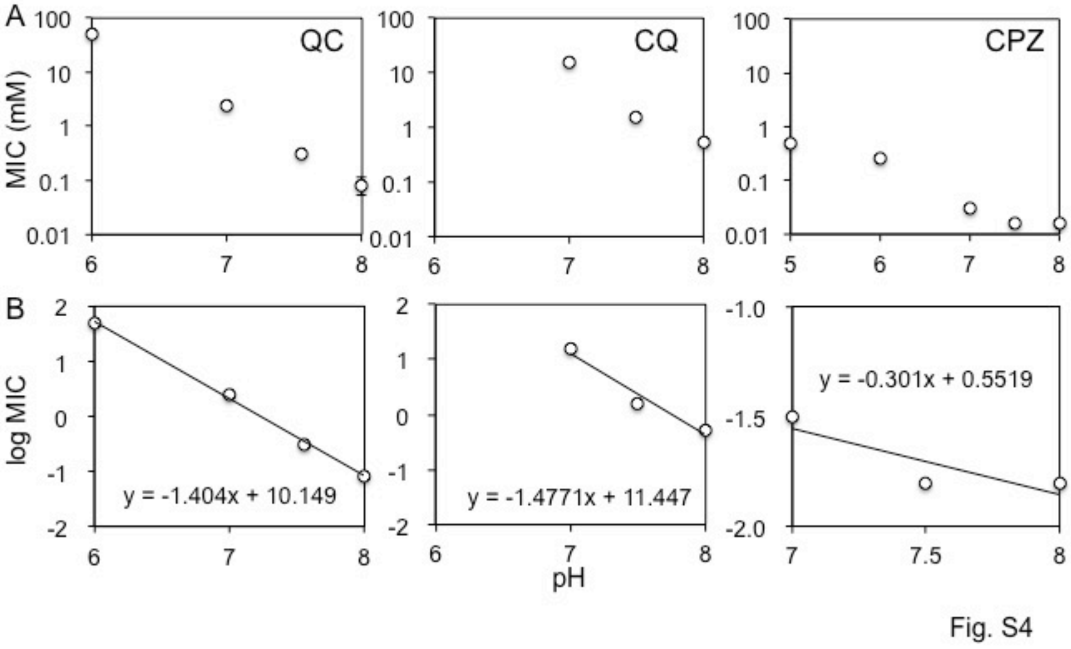


**Fig. S4**. **Linear regression models between the MICs and pH of antimalarial and antipsychotic drugs**

**(A)** The MICs of QC, CQ and CPZ determined in BY4741 cells grown in YPD buffered at each pH value were plotted against the indicated pH value of the culture. CQ had no antimicrobial activity at pH 6, even at the concentration of 100 mM. Data are the mean (SE) ($n \geq 3$). **(B)** The MICs in A were plotted as linear regression models between the log MICs and the pH (7-8) (logD). The equations are indicated in each panel.

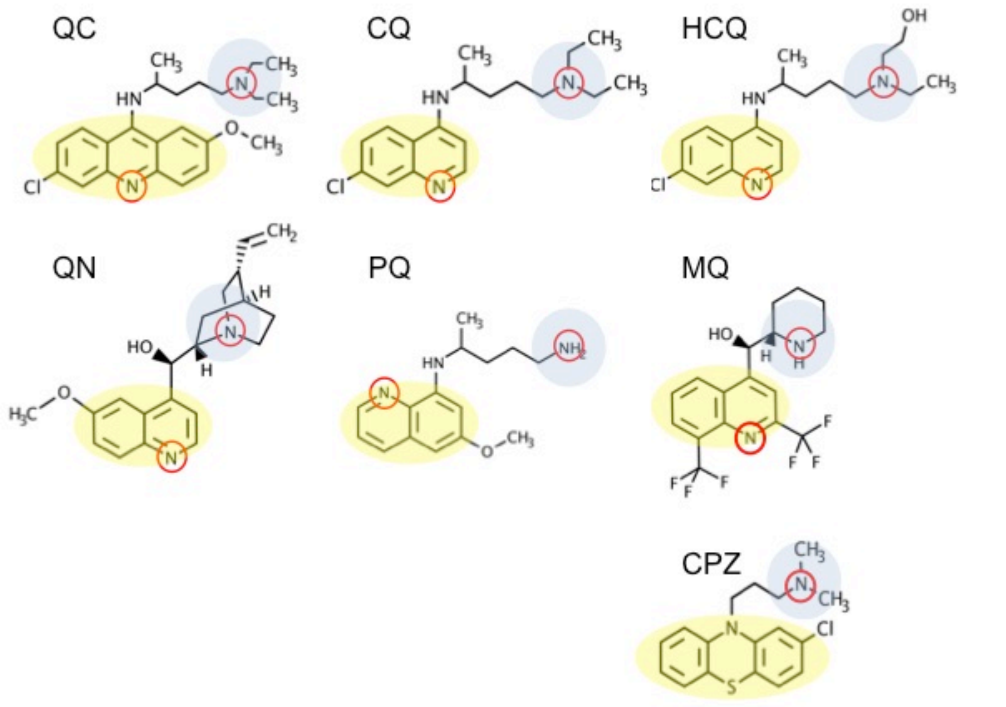


Fig. S5

**Fig. S5. CAD structures of antimalarial drugs and an antipsychotic drug**

Structures of the antimalarial drugs and antipsychotic drug were drawn using MarvinSketch. The protonated nitrogen atoms are enclosed by red circles. Blue and yellow zones indicate hydrophilic and hydrophobic regions, respectively. The abbreviations of the drug names are as follows: QC: quinacrine, CQ: quinacrine, HCQ: hydroxychloroquine, QN: quinine, PQ: primaquine, MQ: mefloquine, and CPZ: chlorpromazine.
